# A molecular basis of somatotopic map formation

**DOI:** 10.1101/2025.08.13.670179

**Authors:** Kevin T. Sangster, Xinying Zhang, Daniel del Toro, Christina Sarantopoulos, Ashley M. Moses, Shreya Mahasenan, Daniel T. Pederick, Sophie Perreault, Catherine Fallet-Bianco, R. Brian Roome, Elena Seiradake, Liqun Luo, Artur Kania

## Abstract

Somatotopy is a recurring organisational feature of the somatosensory system where adjacent neurons and their connections represent adjacent regions of the body. The molecular mechanisms governing the formation of such “body maps” remain largely unknown. Here we demonstrate that the cell surface proteins teneurin-3 and latrophilin-2 are expressed in opposing gradients aligning with the somatotopic map in the dorsal horn of the mouse spinal cord. Genetic manipulation of these proteins in spinal dorsal horn or sensory neurons distorts the somatotopy of neuronal connections and impairs accurate localisation of a noxious stimulus on the surface of the body. Our work provides the foundation for a molecular model of somatotopic map formation and insights into the role of somatotopic maps in the ability to accurately locate somatosensory stimuli or topognosis.

## Introduction

The architecture of neuronal connections reflects their functional logic. A key organisational feature of neuronal circuits are “topographic maps” that maintain the spatial relationship between a group of neurons and their post-synaptic targets. The consequence of this, in the context of sensory processing, is that information concerning adjacent stimuli is communicated by adjacent neurons at multiple relay nodes of a sensory system. The classical paradigm for studying the development of such organisation are retinotopic maps whose connectivity is driven by gradients of cell surface proteins and refined by cellular-level processes^1–7^. However, knowledge of the molecules that wire other sensory maps, particularly those in the somatosensory system, remains limited.

Somatotopic maps are a unique example of topographic maps where nearby neurons represent contiguous anatomical structures. Somatotopic organisation is characteristic of most vertebrate somatosensory circuits, including the dorsal horn of the spinal cord (DH)^8–10^, ventrobasal thalamus^11,12^, and primary somatosensory cortex^13,14^. The rodent DH exhibits a mediolateral (ML) organisation that corresponds to the proximodistal (PD) axis of the body, where the medial DH receives sensory inputs from distal regions (e.g. the paw), while the lateral DH corresponds to more proximal regions of the body (e.g. the shoulder/hip to midline)^8–10^. This organisation is also reflected by neuronal activity within the DH induced by sensory stimuli^15–17^. The ML map is oriented orthogonally to the DH laminae that receive modality-specific sensory inputs^18,19^; the result is a columnar-like architecture where a vertical “columel” processes noxious, thermal, and light touch information originating in a particular location on the surface of the body^16,20^.

Despite the discovery of somatotopic maps over a century ago, the mechanisms that underlie their development remain poorly understood and their precise role in internal representation of our bodies is speculative^13,21–23^. It is known that neuronal activity plays a critical role in the refinement of developing body maps^24–26^. On the other hand, sensory axons are somatotopically sorted immediately upon entry into the DH, suggesting a contribution by hard-wiring molecular signals^27–31^. However, the identity of these has remained elusive: none of the classical neuronal connectivity molecules appear to be involved despite their extensive study. Here, we identify cell surface proteins teneurin-3 (Ten3) and latrophilin-2 (Lphn2) as essential for the formation of the DH somatotopic map and provide evidence that its miswiring attenuates the accuracy of somatosensory stimulus localisation.

### Ten3 and Lphn2 expression reflects the ST organisation of the DH

Recent work has shown that heterophilic repulsion mediated by Ten3 and Lphn2, in combination with Ten3-Ten3 homophilic attraction, establish a point-to-point topography of neuronal connections in the hippocampus^32–34^. To determine whether Lphn2 and Ten3 could direct the development of DH connectivity, we examined their expression in the mouse lumbosacral DH at embryonic day (e) 14.5, when cutaneous afferents begin their entry into spinal laminae I-IV^35,36^. We observed complementary mRNA gradients that spanned the limb somatotopic map: *Lphn2* mRNA expression was high in the medial DH (representing the distal limb) and low in the lateral DH, whereas *Ten3* mRNA expression was high in the lateral DH (representing the proximal limb) and low in the medial DH (Figure 1A). Analysis of protein expression using an anti-Ten3 antibody ^32^ and *Lphn2-mVenus* knock-in mice^37^ revealed similar mediolateral gradients (Figure 1B).

**Figure 1.**
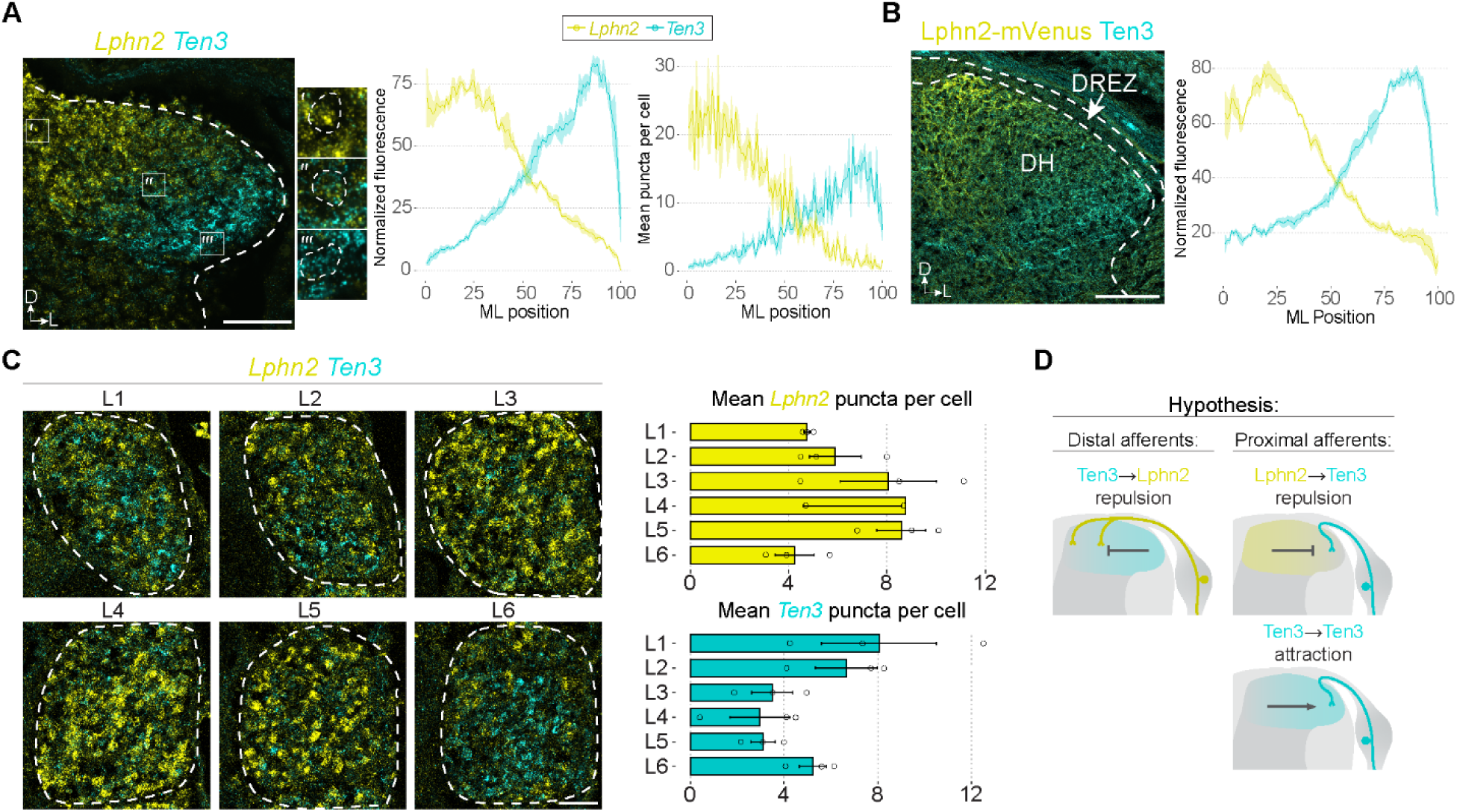
Complementary expression of Ten3 and Lphn2 in the DH and DRG. **A**, *Ten3* and *Lphn2* mRNA localisation in the lumbosacral DH at e14.5 (left). Quantification of RNAscope puncta and fluorescence intensity (right, n = 4 embryos). **b**, Localisation of Ten3 and Lphn2 proteins, with anti-Ten3 and anti-GFP antisera, respectively, in the lumbar DH and adjacent dorsal root entry zone (DREZ) of *Lphn2-mVenus* knock-in mice at e14.5 (left). Quantification of fluorescence intensity in the DH (right, n = 3 embryos) **C**, *Ten3* and *Lphn2* mRNA localisation in lumbar DRG at e14.5 (left) and quantification of mRNA puncta (right, n = 3 embryos). **D**, Hypothesized interactions governing DH somatotopic map formation based on the expression and previously described roles of Ten3 and Lphn2. All data shown as mean ± SEM. Scale bars in (**A**,**B**) = 100 μm, (**C**) = 50 μm. Axis labels in this and subsequent figures: M, medial; L, ventral; D, dorsal; V, ventral.

We then examined the afferent inputs to the DH: the primary sensory neurons residing within the dorsal root ganglia (DRG). Although there is no apparent somatotopic organisation of DRG neurons within individual ganglia, different ganglia innervate distinct regions of skin termed “dermatomes”^8,38,39^. Correspondingly, we did not observe any spatial bias to *Ten3* or *Lphn2* expression in individual ganglia along either the dorsoventral or mediolateral axes (Figure S1A). Meanwhile, we did observe an inverse relationship between *Ten3* and *Lphn2* mRNA levels in individual DRG neurons, where those that expressed high levels of *Ten3* also expressed low levels of *Lphn2* and vice versa (Figure S1B). Additionally, DRG neurons that predominantly express either *Ten3* or *Lphn2* could be identified across all major sensory neuron subtypes (Figure S1C).

Reflecting the dermatome organisation, ganglia enriched in distal hindpaw-innervating neurons (L3-L5) had high *Lphn2* and low *Ten3* levels of mRNA. Conversely, ganglia enriched in more proximally-projecting neurons (L1-2 and L6) had low *Lphn2* and high *Ten3* (Figure 1C). Thus, the sensory neuron expression of *Lphn2* and *Ten3* correlated with DH Lphn2 and Ten3 gradients and the somatotopic map of limb: distal limb sensory neurons that innervate the Lphn2*-*high medial DH express high levels of *Lphn2*, and proximal sensory neurons that innervate the Ten3-high lateral DH express high levels of *Ten3*. A similar correlation could be made between the protein gradients in the DRG axons located in the dorsal root entry zone (DREZ) immediately outside the DH and the gradients in the DH (Figure 1B, quantified in Figure S1D). Next, to determine if Ten3 and Lphn2 gradients align with somatotopic maps in other somatosensory regions, we examined *Ten3* and *Lphn2* mRNA expression in the ventrobasal (VB) thalamus, an important node in the ascending somatosensory pathway, and uncovered a similar gradient that aligned with the VB somatotopic map^11^ (Figure S1E). Finally, to examine whether such Ten3/Lphn2 gradients are evolutionarily conserved we examined the expression of *TENM3* and *ADGRL2* mRNAs, which encode the human homologues of mouse Ten3 and Lphn2, in the human fetal dorsal horn and observed similar mediolateral gradients (Figure S1F). Thus, our observations suggest that Ten3 and Lphn2 are expressed in gradients along the proximodistal axis of somatotopic maps in a conserved manner across the somatosensory system.

### Ten3 and Lphn2 are required for the somatotopic organisation of sensory afferents

Based on the observed expression patterns in the DH and DRG, as well as the previously described functions of these proteins^33^, we hypothesized the following: (1) repulsion from Ten3 in the DH acting on Lphn2 in DRG axons directs distal limb afferents to the medial DH (Figure 1D, left), (2) a combination of repulsion from Lphn2 in the medial DH acting on Ten3 in the DRG axons and attraction from Ten3 in the lateral DH acting on Ten3 in DRG axons cooperate to direct proximal limb afferents to the lateral DH (Figure 1D, right). To test our hypothesis, we first challenged DRG axons with purified Lphn2^Lec^ (the teneurin-binding lectin domain of Lphn2) and the Ten3 extracellular domain. We found that both Lphn2^Lec^ and Ten3 repel DRG axons *in vitro* (Figure S2), thereby demonstrating that Ten3 and Lphn2 can guide DRG axons *in vitro*. Next, we generated conditional knockouts (cKOs) of *Ten3* and *Lphn2* in the DRG and DH (Figure S3A; see methods), hereby referred to as Ten3^ΔDRG^, Lphn2^ΔDRG^, Ten3^ΔDH^, and Lphn2^ΔDH^. To assess the distribution of sensory afferents along the mediolateral extent of the DH, we co-injected cholera toxin subunit B (CTB) and wheat germ agglutinin (WGA) subcutaneously to the hindpaw or dorsal midline. These preferentially label myelinated (e.g. innocuous touch-relaying) and unmyelinated (e.g. pain and temperature-relaying) sensory afferents, respectively^10,26,40^.

First, we examined distal limb afferent inputs to the DH, which we hypothesized to be guided by Ten3 (DH) → Lphn2 (DRG) repulsion (Figure 1D, left). We examined these afferents in Ten3^ΔDH^ and Lphn2^ΔDRG^ cKO mice and littermate controls by injecting CTB/WGA into the hindpaw and quantified the distribution of the labelled axons along the ML and superficial-deep (SD) axes of the DH (Figure S3B; see methods). In wild-type mice, the paw representation is disproportionally large and duplicated at the most medial extreme of the dorsal horn^10,16,26,41^. In our controls, we saw the expected wild-type distribution of afferents, however in Ten3^ΔDH^ cKO mice we observed a dramatic shift in the ML distribution of labelled afferents (Figures 2A-C and S3C). Despite the shift in ML position, CTB and WGA-labelled axons remained closely aligned with each other along the ML axis (Figure S3D), and there was no shift in the distribution of CTB and WGA-labelled axons along the orthogonal SD axis (Figures S3E-S3G). The Lphn2^ΔDRG^ cKO mice exhibited a milder but significant shift along the ML axis and no change in the SD axis distribution or in the ML alignment between CTB and WGA-labelled afferents (Figures 2D-F and S3H-L).

**Figure 2.**
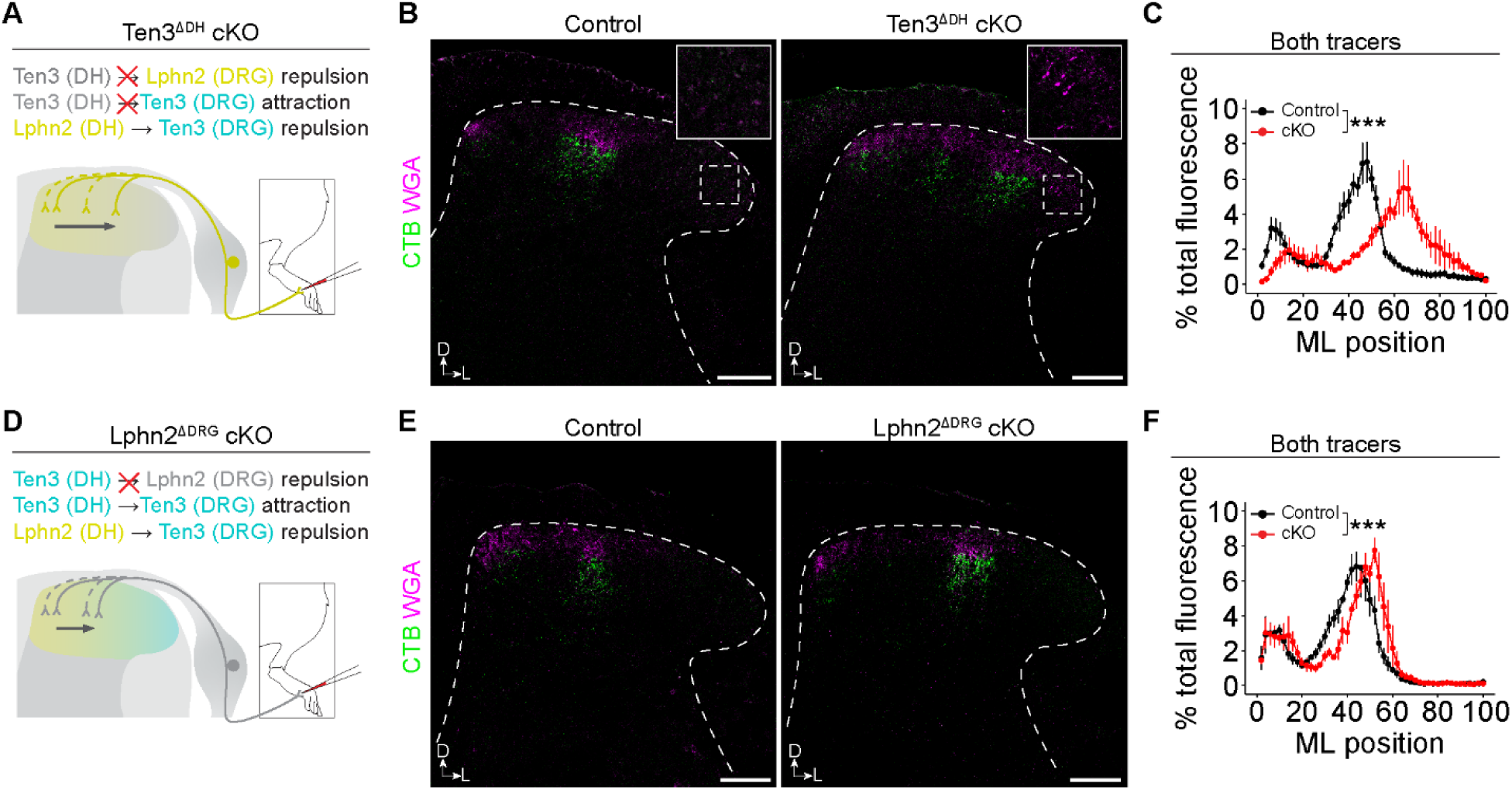
Aberrant somatotopy of distal sensory afferents in the DH of Ten3 and Lphn2 cKOs. **A**,**D**, Schematic of lost and preserved interactions in (**A**) Ten3^ΔDH^ cKOs and (**D**) Lphn2^ΔDRG^ cKOs (top) and the observed shift in the ST distribution of sensory afferents in the cKOs (bottom, filled lines) compared to controls (bottom, dotted lines). **B**,**E**, Representative images of CTB and WGA labeled afferent termini in (**B**) Ten3^ΔDH^ cKOs (insets showing WGA near lateral edge of DH in cKOs) and (**E**) Lphn2^ΔDRG^ cKOs compared to controls. Stippled lines outline the DH. **C**,**F**, Quantification of the distribution of CTB and WGA signal across the ML axis of the DH in cKOs and control littermates in (**C**) Ten3^ΔDH^ cKOs (n = 4 controls, 4 cKOs) and (**F**) Lphn2^ΔDRG^ cKOs (n = 4 controls, 3 cKOs). All data shown as mean ± SEM. Two-way mixed ANOVA was performed and genotype x bin interaction terms are reported. ***P ≤ 0.001; **P ≤ 0.01; *P ≤ 0.05. Scale bars = 100 μm.

Next, we examined proximal afferent inputs to the DH, which we hypothesized to be guided by a combination of Lphn2 (DH) → Ten3 (DRG) repulsion and Ten3 (DH) → Ten3 (DRG) attraction (Figure 1D, right). We therefore examined these afferents in Ten3^ΔDH^, Lphn2^ΔDH^, and Ten3^ΔDRG^ cKO mice and compared them to littermate controls by injecting CTB/WGA subcutaneously at the dorsal midline. Like for the distal limb afferents, we observed a dramatic shift in the ML distribution of proximal afferents in the Ten3^ΔDH^ cKO mice (Figures 3A-C and S4A). However, this was accompanied by a decrease in the ML alignment of the CTB with the WGA-labelled afferents (Figure S4B), and an expansion in the SD distribution of both classes of afferents (Figures S4C-S4E). Lphn2^ΔDH^ cKO mice exhibited a comparatively mild but significant shift along the ML and SD axes, without a consequence on the alignment of CTB with the WGA-labelled afferents (Figures 3D-F and S4F-J). Unexpectedly, the Ten3^ΔDRG^ cKO mice did not exhibit any significant changes in the targeting of afferents in the DH (Figures 3G-I and S4K-O), indicating that Ten3 in the DRG may be dispensable for correct somatotopic and laminar targeting. Importantly, Ten3 (DH) → Lphn2 (DRG) repulsion is likely intact in Ten3^ΔDRG^ cKO mice, suggesting that this repulsion alone may be sufficient to separate the distal and proximal inputs along the ML axis. Together, our cKO experiments demonstrate that Ten3 and Lphn2 are essential for the correct somatotopic organisation of primary sensory afferents in the DH, with the severity of miswiring phenotypes depending on the gene deleted and the targeted tissue.

**Figure 3.**
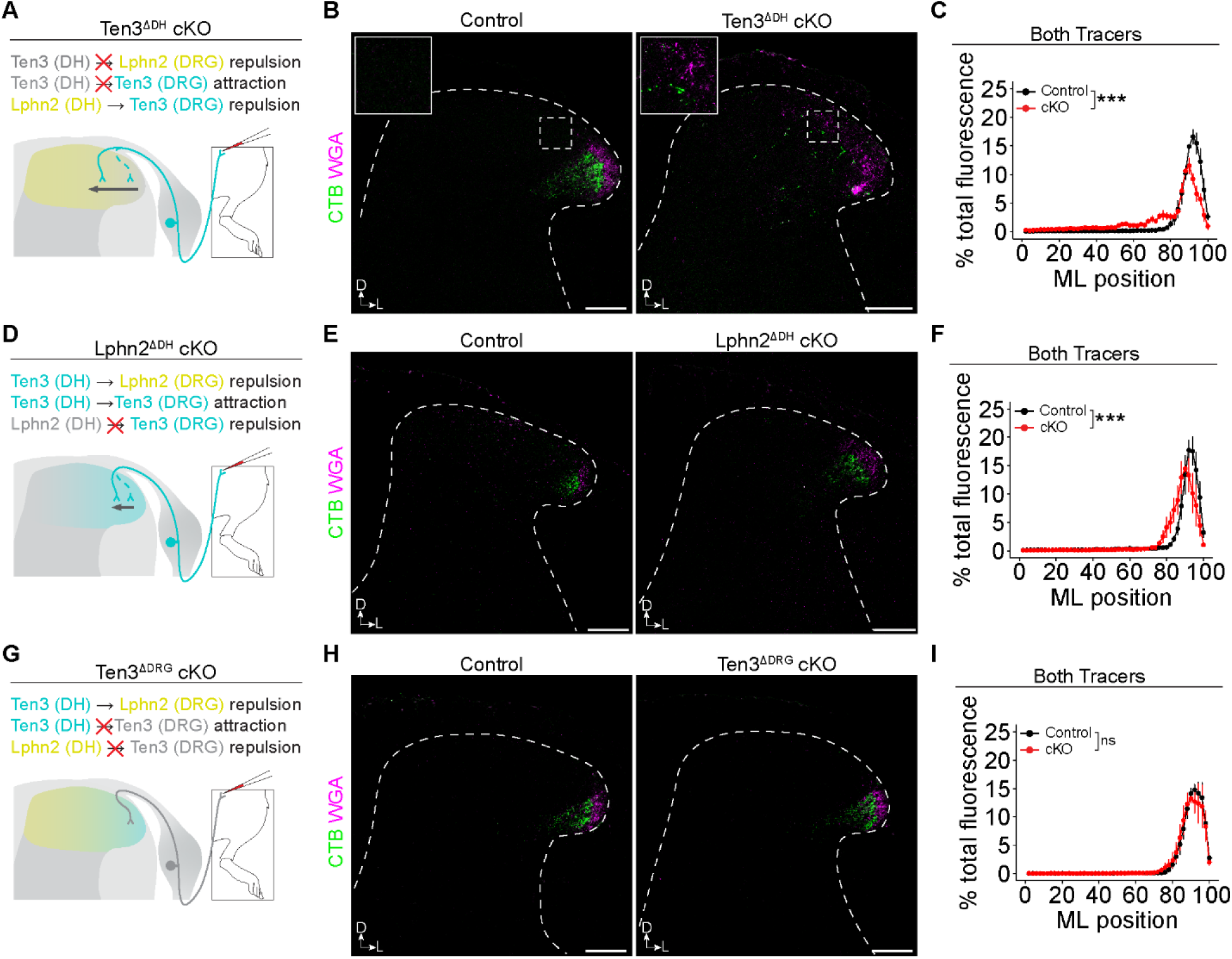
Altered somatotopy of proximal sensory afferents in the DH of Ten3 and Lphn2 cKOs. **A**, **D**, and **G**, Schematic of lost and preserved interactions in (**A**) Ten3^ΔDH^ cKOs, (**D**) Lphn2^ΔDH^ cKOs, and (**g**) Ten3^ΔDRG^ cKOs (top) and the observed shift in the ST distribution of afferent termini in the DH of cKOs (bottom, filled lines) compared to controls (bottom, dotted lines). **B**, **E**, and **H**, Representative images of CTB and WGA labeled afferents in (**B**) Ten3^ΔDH^ cKOs (insets show more medially located CTB/WGA in cKOs), (**E**) Lphn2^ΔDH^ cKOs, and (**H**) Ten3^ΔDRG^ cKOs compared to controls. Stippled lines outline the DH. **C**, **F**, and **I**, Quantification of the distribution of CTB and WGA signal across the ML axis of the DH in cKOs and control littermates in (**C**) Ten3^ΔDH^ cKOs, (**F**) Lphn2^ΔDH^ cKOs, and (**I**) Ten3^ΔDRG^ cKOs (n = 4 controls and 4 cKOs for all). All data shown as mean ± SEM. Two-way mixed ANOVA was performed and genotype x bin interaction terms are reported. ***P ≤ 0.001; **P ≤ 0.01; *P ≤ 0.05. Scale bars = 100 μm.

### Computational modelling of somatotopic map formation in the DH

To interpret the observed phenotypes and gain further insights into the developmental programs that produce the DH somatotopic map, we constructed a comprehensive computational model of DH somatotopic map formation. Over the past 40 years, several computational models of topographic map formation in the visual system have been developed and iteratively improved. We first determined if these theoretical models of topographic map formation could be applied to somatotopic maps. We therefore adapted the Koulakov model^7,42,43^ (see methods) as it is a relatively abstract model without excessive constraints regarding the cellular mechanisms. It also effectively predicts the observed effects of genetic manipulations^44^, and incorporates guidance cues, competition, and activity-dependent refinement into a single unified model^7^. After adapting this model to the DH somatotopic map, we then performed Bayesian Simulation-Based Inference (SBI) using the Sequential Monte Carlo method for Approximate Bayesian Computation (ABC-SMC) to identify parameter values for our model that fit both the observed control and cKO data in an unbiased manner (Figure 4A; see methods).

**Figure 4.**
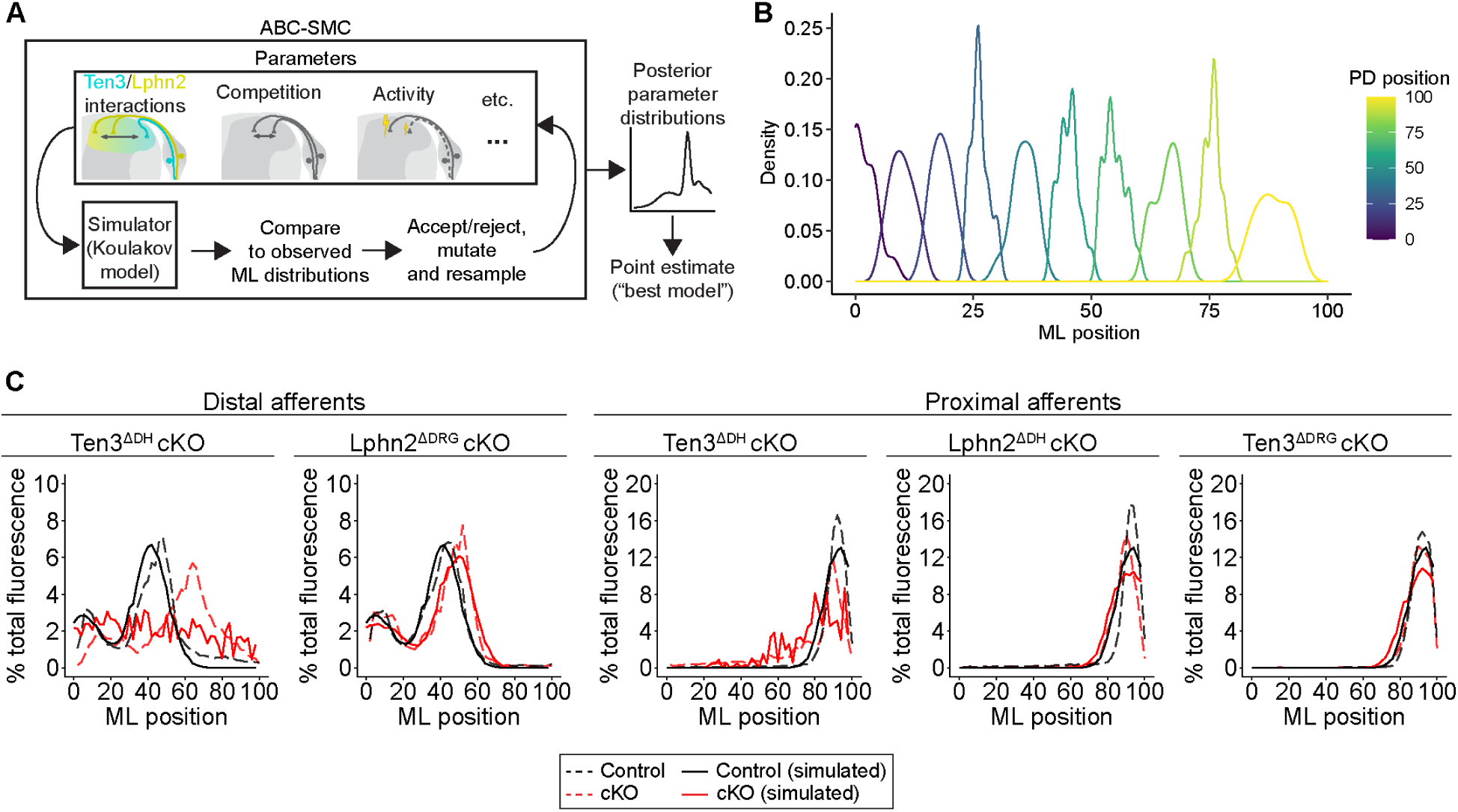
Computational modelling of DH somatotopic map formation. **A**, Simplified schematic of model parameters and of the ABC-SMC procedure. **B**, Correct distal → medial and proximal → lateral mapping from one representative simulation under wild-type conditions. Simulated ML density of terminal zones in the DH from 10 DRG axons selected across the PD axis. **C**, Observed and simulated tracer injections for controls and cKOs across all genotype/injection site combinations examined. Average distributions of each condition are shown. Simulated distribution taken from the average of 10 independent simulations.

Using the parameter values fit above, we first confirmed that this model was able to accurately reproduce the proximal → lateral and distal → medial connectivity between DRG and DH neurons under wild-type conditions (Figures 4B and S5A). We then confirmed that we were able to closely recapitulate the observed distributions of CTB/WGA along the ML axis in both controls and all cKOs examined experimentally above (Figure 4C). Inspection of the parameter values (table S1) revealed that the best fitting model incorporates a prominent role for activity-dependent refinement of connectivity, consistent with previous observations^24–26^. Furthermore, disrupting activity dependent refinement in our model produced a broadening of DRG axon terminal zones and disruptions to DH somatotopy that was qualitatively similar to experimental observations (Figures S5B-D), validating this component of the model. Our model also indicates that there is strong competition between DRG axons for DH space along the ML axis, consistent with previous results showing an expansion of the saphenous nerve afferent termini when the afferents of the sciatic nerve are eliminated^45,46^. We recreated this experiment *in silico* by ablating proximal limb DRG neurons and observed a similar expansion of the remaining axons to fill the available ML extent of the DH (Figures S5E and S5F), further validating our model. Additional inferences of our model are that Ten3 (DH) → Lphn2 (DRG) repulsion plays a more prominent role than Lphn2 (DH) → Ten3 (DRG) repulsion or Ten3 (DH) → Ten3 (DRG) attraction, and that the loss of Lphn2 or Ten3 likely impacts both guidance and activity-dependent refinement, particularly in the Ten3^ΔDH^ cKOs. Taken together, we show that despite being developed for a different sensory system patterned by gradients of unrelated molecular cues, a theoretical model of topographic map formation in the visual system can be adapted to accurately represent the developing somatotopic map in the DH.

### Ten3^ΔDH^ cKOs exhibit an altered somatotopic distribution of neural activity and behavioural responses

Given the observed miswiring phenotypes, we asked if this miswiring was accompanied by alterations in the function of the DH neuronal circuitry. Towards this end, we selected the cKO with the most severe miswiring phenotype (Ten3^ΔDH^) and examined the consequence of injecting the hindpaw with formalin, a noxious stimulus that results in a somatotopically restricted induction of the neuronal activity marker c-Fos in DH neurons^15^. Following injection of formalin into the distal hindpaw, both cKO mice and control littermates displayed a significant induction of c-Fos in the DH ipsilateral to the injection (Figures S6A and S6B). While controls showed increased c-Fos expression predominantly in the somatotopically appropriate medial DH, Ten3^ΔDH^ cKO mice exhibited a greater number of c-Fos-expressing cells in the lateral DH (Figures 5A-D). Matching the more widespread induction of c-Fos, cKO mice also had more c-Fos-expressing cells on the side ipsilateral to the injection when compared to control littermates (Figures S6C-S6E).

**Figure 5.**
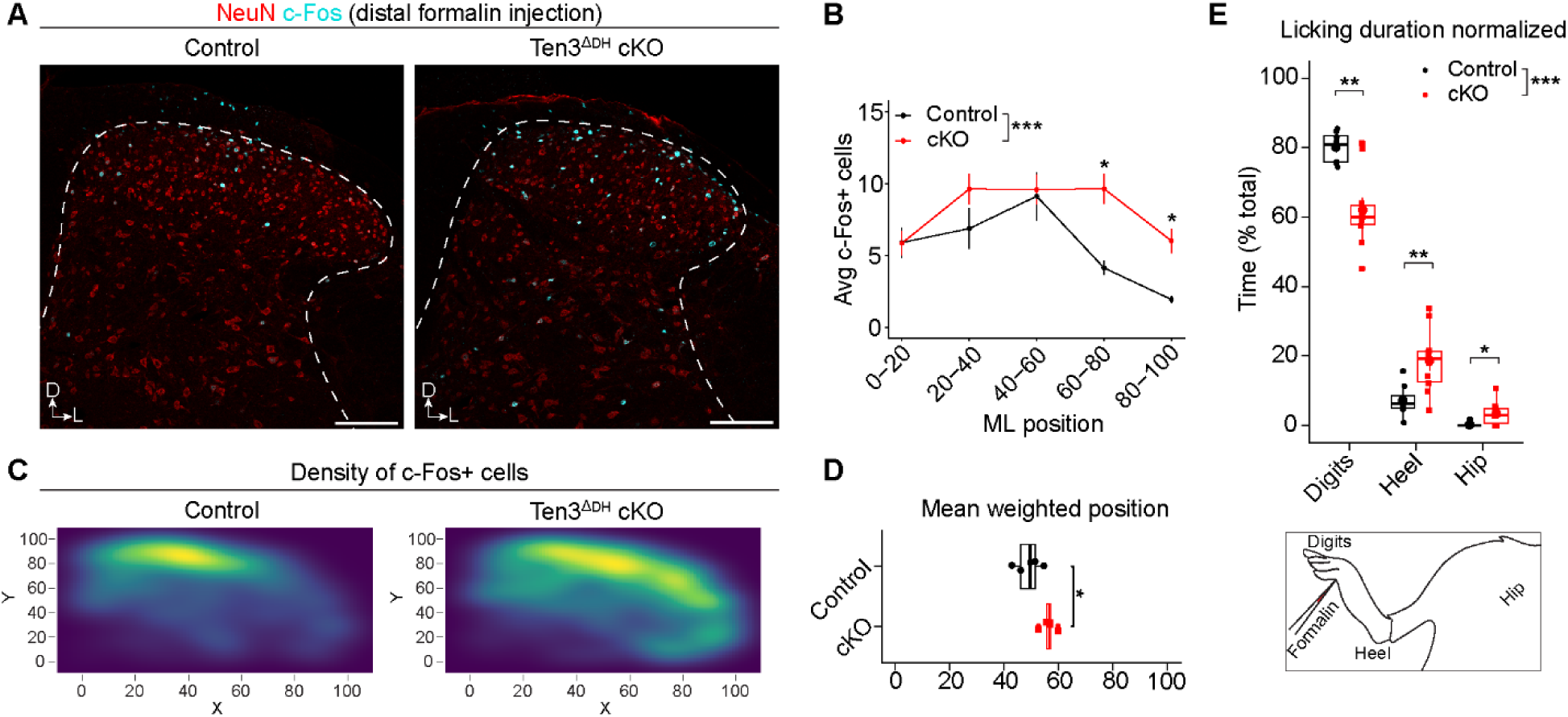
Altered distribution of c-Fos expression in the DH and behavioural responses induced by a noxious stimulus in Ten3^ΔDH^ cKOs. **A**, Representative images of c-Fos distribution in the DH induced by formalin injection to the hindpaw of Ten3^ΔDH^ cKOs and contols. **B-D**, Quantification of c-Fos-expressing cells in Ten3^ΔDH^ cKO mice and control littermates (n = 5 controls and 5 cKOs). (**B**) Distribution of c-Fos positive cells across the ML axis of the DH. Two-way mixed ANOVA test was performed, and genotype x bin interaction term is reported along with Benjamini-Yekutieli corrected post-hoc p-values for individual bin comparisons. (**C**) 2D kernel density plots showing their distribution of Fos positive cells in the DH. Hotelling T^2^ p-value 0.03. (**D**) Average ML position of c-Fos-expressing cells compared with Welch’s t-test. **E**, Quantification of nocifensive licking behaviour in response to formalin injection to the hindpaw near the digits, depicted as a percent of total linking time per limb region (n = 8 controls and 10 cKOs). Two-way mixed ANOVA was performed and genotype x location interaction term is reported. Post-hoc Mann-Whitney tests were performed for each limb location. All data shown as mean ± SEM. ***P ≤ 0.001; **P ≤ 0.01; *P ≤ 0.05. Scale bars = 100 μm.

In addition to being able to induce somatotopically restricted c-Fos activity, formalin injection elicits highly accurate licking of the injection site. Thus, using this paradigm we sought to examine the functional impact of somatotopic miswiring on the ability to locate a stimulus on the surface of the body, or topognosis. We first examined the precision of topognosis in Ten3 constitutive knockout (Ten3^KO^) mice^32,47^. Compared to control littermates, Ten3^KO^ mice spent less time licking regions of the limb close to the injection site and more time licking regions of the limb farther from the injection (Figures 6F and G), potentially indicating a deficit in their ability to precisely locate the stimulus. Meanwhile, the total duration of licking, an indication of stimulus intensity, was not different between KO and control mice (Figure 6H). However, Ten3^KO^ mice also showed defects in the ability to remove an adhesive tape placed on their hindlimbs, suggesting broad deficits in sensorimotor coordination (Figures S6I and S6J).

We next assessed topognosis in Ten3^ΔDH^ cKO mice given the magnitude of their anatomical miswiring phenotype. Similar to the Ten3^KO^ mice, Ten3^ΔDH^ cKO mice exhibited a reduction in time spent licking areas close to the injection and increase in time licking areas farther from the injection site (Figures 5E and S6K), indicating impaired topognosis. This phenotype was not accompanied by any significant changes in total licking duration (Figure S6L). Finally, unlike the Ten3^KO^ mice, Ten3^ΔDH^ mice performed normally in the adhesive removal assay (Figures S6M and S6N), indicating normal sensorimotor coordination. Together, these data indicate that loss of Ten3 in the DH, in addition to causing significant disruptions to DH somatotopy, impairs somatosensory topognosis.

## Discussion

Our study provides foundational insights into the molecular logic of somatotopic map formation. First, our experimental observations and modelling argue that the somatotopic map formed between peripheral sensory neurons and those in the spinal DH is driven by graded expression of Ten3 and Lphn2. This complements previous studies on their role in establishing of neuronal connections through binary or all-or-nothing attractive and repulsive signalling. Additionally, we also provide direct functional evidence of the importance of somatotopic maps in the ability to localise a stimulus on the surface of the body.

Several parallels and distinctions between our work and that on the formation of retinotopic maps can be made. Like in retinotopic maps, past studies and this work suggest that axon-axon interactions such as pre-ordering prior to arrival in target area and inter-axon competition, as well as activity-dependent refinement, play an important role in the establishment of the DH somatotopic map. However, while disrupting EphA/ephrin-A signalling largely results in the formation of additional discrete terminal zones of retinal axons in the superior colliculus^5^, we generally observe broadening and/or shifting of sensory axon terminal zones in the ML axis of the DH, without the formation of additional ones, suggesting fundamental cellular/molecular differences between these processes. Of particular interest would be the significance of Ten3/Lphn2 and EphA/ephrin-A co-expression, the latter of which have been implicated in somatosensory cortex somatotopic map formation by specifying afferent thalamic axon connectivity^48,49^. Finally, the presence of Ten3/Lphn2 gradients in many regions of the developing nervous system raises the possibility that they pattern all somatotopic maps within the somatosensory system, and more generally, circuits whose organising principle requires the preservation of spatial relationship between neurons and their targets^50^.

Although we cannot rule out other DH functions of Ten3 contributing to the impaired topognosis phenotype, to our knowledge, our data represent the only functional evidence indicating a requirement for somatotopic maps in topognosis. More generally, our data adds to the currently limited behavioural evidence that topographic maps are required for correct sensory processing. Finally, the unchanged duration of licking in response to formalin in Ten3^ΔDH^ cKOs suggests that stimulus intensity and location can be dissociated, echoing the idea of parallel motivational/affective and discriminatory systems underlying nociception^51^. Our genetics-based precise manipulation of a somatotopic map provides a new inroad into questions such as body map plasticity, the integration of somatotopic somatosensory maps with somatotopic motor maps, or those of other sensory modalities, and their overall function in representing our bodies and whole-body action planning^23^.

## Methods

### Acquisition of human tissue

Collection of the human fetal spinal cord samples were obtained with the parents’ written informed consent through a spontaneous or induced abortion donor program at the Centre hospitalier universitaire - Sainte-Justine (CHU-SJ), approved by the IRCM and CHU-SJ Research Ethics Committee under the name “Pour une meilleure compréhension du développement de la moelle épinière et des maladies qui y sont associées”. Donor spinal cords were dissected and fixed for 24h in formalin, then processed for cryosectioning as below.

### Mouse lines

All animal studies were approved by the IRCM’s Animal Care Committee in agreement with Canadian Council on Animal Care (CCAC) guidelines, as per protocol numbers 2021-12AK and 2023-11AK. Mice were group-housed with access to food and water *ad libitum*. All mice were maintained on a mixed C57BL/6 and 129/Sv background and both male and female mice were used in all experiments. Mouse lines used in this study include *Lbx1:Cre*^52^*, PLAT:Cre*^53^*, Ten3^fl(^*^32)^, *Lphn2^fl(^*^37)^*, Lphn2-mVenus*^37^, and *Ten3^KO^*^(47)^. Within the spinal cord, *Lbx1:Cre* drives recombination within the dI4/dILA, dI5/dILB neurons which make up the majority of the DH while sparing most other spinal populations and the DRG^54–56^. *PLAT:cre* is expressed in neural crest derived cells, including DRG neurons, while sparing the spinal cord and brain^53^. *PLAT:Cre* was chosen over alternatives like *Pirt:Cre* and *Advilin:Cre* since these are expressed in neural crest-derived cells as early as Theiler Stage 15 (e9.5) and throughout DRG neurons by e11.5 (Figure S2A).

### Cutaneous CTB/WGA injections

Adult mice (>8 weeks old) were anaesthetised via continuous inhalation of isofluorane (1-3%) and injected with 4 µL of a mixture of 5 mg/mL CTB-AF488 (Thermo Fisher Scientific, cat. #C34775) and 20 mg/mL WGA-AF555 (Thermo Fisher Scientific, cat. #W32464) dissolved in sterile PBS. Blunt forceps were used to pinch and stabilize the skin and injections were performed into the dermis using a beveled glass capillary needle. Distal injections were performed to the glabrous skin of the hindpaw between the two distal-most promontories. Proximal injections were performed at the dorsal midline at the same rostrocaudal level as the iliac crest. All mice were perfused after three days following the injection as described below.

### Adhesive removal assay

The adhesive removal assay was performed as previously described^57,58^. In brief, mice were tested on their ability to remove an adhesive (half of a 1.5 mL Eppendorf tube cap label, cut into a semicircle) placed on the plantar surface of the hindpaw. The assay was repeated for five consecutive days, with the first four days considered training days and only the final day used for comparison. The latency to remove the adhesive was recorded in seconds, with a maximum time of 30 minutes. If mice exceeded the maximum time, it was marked as a “failure to remove” and the latency was recorded as 30 minutes for the purpose of data analysis. Mice then returned to their home cage.

### Formalin Assay and c-Fos induction

The formalin assay was performed as previously described^58–60^. Mice were first habituated to the testing chamber (a clear plastic cylinder filmed from below). They were then subcutaneously injected with 20 μL of 2% formalin (in 0.9% saline) to the plantar surface of the hind paw using a 28G needle and returned to the chamber. Mice were filmed from below for 60 minutes following injection. All quantification of time spent licking each region of the body was done in the acute phase (0-10 minutes after injection). Mice were then anesthetized and, after an additional 30 minutes, perfused as described below to analyze the distribution of c-Fos protein (after a total of 90 minutes following the stimulus).

### Tissue fixation, freezing and sectioning

Mice were anesthetized via intraperitoneal injection of a Ketamine/Xylazine mixture (0.1mL/g body weight of 10 mg/mL Ketamine, 1 mg/mL Xyaline, in 0.9% saline). Mice were then perfused with approximately 15 mL of 1x PBS followed by 15 mL of 4% PFA in PBS. Spinal cords were post-fixed in 4% PFA overnight, washed 3x with PBS and left in a 30% sucrose in PBS solution until they had sunk. Tissue was cryosectioned into 25 µm sections for adult tissue used for immunohistochemistry and 12 µm sections for embryonic tissue and adult tissue used for mRNA detection using RNAscope (Advanced Cell Diagnostics (ACDbio)).

### DRG explant culture on Ten3/Lphn2 stripes

DRG explants from E15.5 embryos were dissected and plated on 6 cm dishes containing stripes, prepared essentially as previously described^61^. Briefly, 100 μg/mL of Fc recombinant protein, Lphn2-Lec or Tenm3, were mixed with Alexa 594-conjugated anti-hFc antibody (Thermo Fisher Scientific, cat. #A-11014) in PBS. Proteins were injected into matrices (90 μm width) and placed on 60 mm dishes, resulting in red fluorescent stripes. After 30 min incubation at 37°C, dishes were washed with PBS and matrices removed. Dishes were coated with 100 μg/mL Fc conjugated with anti-hFc (Jackson ImmunoResearch, cat. #109-001-008) for 30 min at 37°C and washed with PBS. Stripes were further coated with 20 μg/mL Laminin in PBS for at least 2 h and washed with PBS. Explants were then cultured for 1-2 days in vitro at 37°C, 5% CO2 in Neurobasal medium plus B27 supplement with the addition of 20 ng/mL Nerve Growth Factor (NGF) and 10 ng/mL Glial cell line-Derived Neurotrophic Factor (GDNF). Samples were then fixed with 4% paraformaldehyde for 20 minutes and stained with rabbit monoclonal anti-beta-III tubulin antibody (Sigma-Aldrich, cat. #ZRB1140) after 20 min permeabilization in 1% BSA, 0.1% Triton X-100 in PBS. Cy2 anti-rabbit IgG secondary antibody (Jackson ImmunoResearch, cat. #111-225-144) was used to detect tubulin. For all conditions n ≥ 15 DRG explants were used. The numbers of beta-III-tubulin-positive (green) pixels on red or black stripes were quantified with ImageJ (version 1.54p)^62^ using a custom-made automatic macro (available upon request).

### Immunohistochemistry

Tissue sections were washed three times with 1x PBS for 5 minutes, blocked for 30 minutes using a solution of 5% heat-inactivated horse serum (HIHS) and 0.1% Triton X-100 in PBS (0.1% tPBS). They were then incubated primary antibody solution (primary antibody in 1% HIHS, 0.1% tPBS) overnight and washed three times with 1x PBS for 5 minutes before being incubated with a secondary antibody (in 1% HIHS, 0.1% tPBS) for 1 hour at room temperature, washed three more times with 1x PBS for 5 minutes, and counterstained with DAPI (4′,6-diamidino-2-phenylindole, Fisher Scientific, cat. # D1306). Slides were coverslipped using a solution of 10% Mowiol (Sigma) and 25% glycerol, then allowed to dry prior to imaging. Primary antibodies used were Rabbit anti-Ten3^32^ (1:500), Goat anti-GFP (1:500, Abcam, ab6673), Mouse anti-NeuN (1:1000, Millipore Sigma, MAB377), Rabbit anti-WGA (1:500, Millipore Sigma, T4144), Goat anti-CTB (1:5000, List Biological Laboratories, #703), Rabbit anti-c-Fos (1:1000, Cell Signaling Technology, #2250S), Goat anti-TrkA (1:500, R&D Systems, AF1056), Goat anti-TrkB (1:500, R&D Systems, AF1494), and Goat anti-TrkC (1:500, R&D Systems, AF373). Donkey secondary antibodies conjugated to Alexa 488, Cy3, or Cy5 were used (1:500, Jackson ImmunoResearch). Micrographs of tissue sections were acquired using the Leica DM6 or SP8 microscopes.

### RNAscope ISH and analysis

RNAscope *in situ* mRNA detection was performed according to the manufacturer’s protocol using the RNAscope multiplex fluorescent reagent kit (ACDbio, cat. #323100). Probes used were Mm-Tenm3-C1 (ACDbio, cat. #411951) and Mm-Lphn2-C2 (ACDbio, cat. #319341-C2). All images of RNAscope-stained sections were acquired using a Leica SP8 confocal microscope. RNAscope images were quantified using QuPath bioimage analysis software (version 0.4.4) ^63^. Nuclear segmentation was performed using StarDist^64^ and individual puncti and clusters were identified using the QuPath subcellular object detection pipeline. Subcellular detection counts associated with individual cells and their associated coordinates were exported from QuPath and analyzed further in R^65^ (version 4.0.5). The x coordinate location of each cell (mediolateral position) was remapped to a 100-unit standard space and counts for a cells within a given area were binned and visualized as mean counts per cell ± one standard error.

### CTB and WGA Image analysis

Initial preprocessing of images from CTB/WGA injections was performed using Fiji^66^ (version 2.14.0/1.54f). Images were first cropped to the dorsal horn up until lamina V and an outline was manually drawn to indicate the region of interest (ROI). For analyses of both tracers, the two channels were first merged using the imageCalculator “Add” function. Fluorescence intensity from all channels were scaled using “Enhance Contrast” with “saturated=0.1” before being thresholded using the auto threshold method “MaxEntropy dark”. Preprocessed images and manually drawn ROIs were then further analyzed in R^65^ (version 4.0.5). To account for the curvature of the dorsal horn (with laminae curved downward almost 90 degrees at the lateral extreme), pixel coordinates within each ROI were remapped to a set of anatomical coordinates according to the following functions:

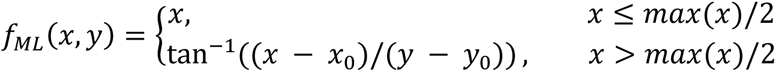

for mapping *x* and *y* pixel coordinates to ML coordinates, and

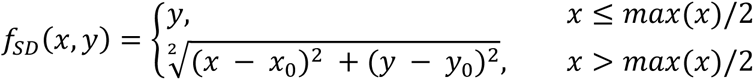

for mapping *x* and *y* pixel coordinates to SD coordinates. Where (*x*_0_, *y*_0_) is the origin, with x_0_ set at the midpoint of the ROI and y_0_ set at the lamina IV/V border for *f_SD_* and the lamina V/VI border for *f_ML_*. *Max*(*x*) refers to the maximum *x* coordinate within the ROI and not the whole image. ML and SD coordinates were then normalized to between 0 and 1. Representative examples of these remapped coordinates are provided here (Figure S2B). The DH was then divided into 50 bins along the ML or SD axes and the fluorescence intensity in each bin is represented as a percent of the total signal (such that the values of all bins sum to 100). Since the bins are created based on distance along the given axis, they are not necessarily of equal size (also given the variability of DH anatomy) and therefore the fluorescence intensity is additionally normalized to bin area.

### c-Fos image analysis

Counts of c-Fos-positive cells was performed manually using the Cell Counter plugin for Fiji^66^ (version 2.14.0/1.54f) and xml files containing the coordinates of each marker were exported for further processing in R^65^ (version 4.0.5). Marker coordinates were remapped to standardized anatomical coordinates as described above. For the remainder of the analyses, the DH was then divided into 5 bins along the ML or SD axes and the raw unnormalized count of c-Fos positive cells per bin was used.

### Computational modelling

To adapt the Koulakov model^7,42,43^ to the DH ST map, we added parameters governing the strength of Ten3/Lphn2 repulsion/attraction since the original model assumed all repulsive/attractive interactions to be of equal strength which may not hold in our case. We also reasoned that it was possible for activity-dependent refinement to be disrupted in our cKOs, either due to the synaptic roles of Teneurins and Latrophilins^67^, or as a secondary circuit-level consequence of ST miswiring, which might then further contribute to the observed ML distribution shifts. We thus incorporated into our model the ability of cKOs to affect activity-dependent refinement and added additional parameters to govern how this interaction occurs, if at all. These parameters determine how much the effect of activity-dependent refinement should be dampened in each cell based on the decrease in Ten3 or Lphn2 in that cell compared to the amount it would express under wild-type conditions using the following formula:

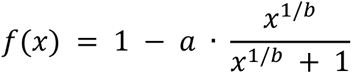

Where x is the absolute decrease in expression compared to wild-type conditions for that cell. For a complete list of model parameters and their description see table S1. The Koulakov model stochastically adds or removes connections to attempt to minimize a global energy value *E* = *E_chem_* + *E_act_* + *E_comp_* that contains three components: chemoaffinity energy (*E_chem_*), activity energy (*E_act_*), and competition energy (*E_comp_*) where the probability of adding or removing a connection proportional to the change in energy. *E_chem_* is calculated according to the following equation:

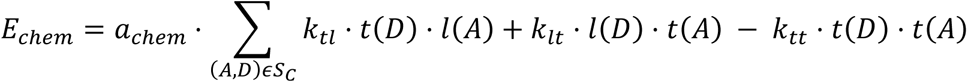

With functions *t* and *l* describing the levels of Ten3 and Lphn2 in a given axon *A* or dendrite *D* and *S_C_* denoting the set of all connected pairs of axons and dendrites. Meanwhile *E_act_* is calculated according to:

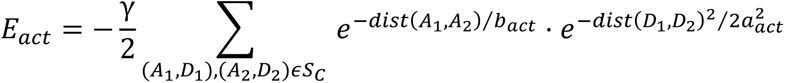

and *E_comp_* by:

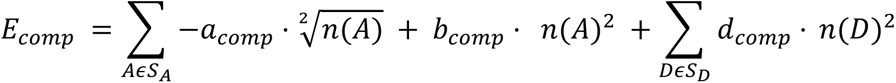

Where *n* describes the number of synapses formed by a given axon or dendrite and *S_A_* and *S_D_* describing the sets of all axons and dendrites respectively. In all simulations we used a linear array of 50 DH neurons across the ML axis with a matching number of DRG axons and ran the simulation for 25,000 iterations (500 steps per axon), since we found the energy *E* for most combinations of parameters to plateau by this point. To simulate the tracer injections, each DRG axon was given an amount of tracer according to a gaussian distribution centered at a particular PD location with the center, σ (determining the PD spread of the tracer), and concentration (determining the total amount of tracer per axon) being determined by simple grid search to best match the observed tracer distributions each ABC-SMC iteration. For simulated distal injections, two gaussians were chosen to mimic the duplicated medial representation of the distal limb. Parameter values for the Koulakov model were fit using ABC-SMC^68^ using the PyABC package (version 0.12.16)^69^. Uniform priors were used for all parameter values. PyABC was run until the “epsilon over walltime” value began to plateau, with a population size of 1000, a maximum number of populations equal to 10, and epsilon values determined using the quantile epsilon method with the parameter alpha equal to 0.1. From this, we obtained posterior distributions over the simulation parameters and a point estimate for the best model (table S1). Simulations of “nerve cut” experiments^45,46^ were run using the parameters from our best fit model and eliminating the lateral half of axons innervating the DH. Simulations of experiments disrupting activity-dependent refinement^24,26^ were run using the parameters from our best fit model with an 80% reduction in the gamma value.

## Statistical Analysis

All statistical analyses were performed in R^65^ (version 4.0.5) and the specific statistical tests used are described in figure legends. Figures generated in R using ggplot2^70^ (version 3.4.2). The two-way mixed ANOVA tests were performed using the rstatix package^71^ (version 0.7.1) according to the formula: fluorescence ∼ genotype*position + Error(ID/position) and the genotype*position interaction terms are reported.

## Data and code availability

Code is available at (https://github.com/k-sangster/somatotopic_maps). All other data are available in the main paper and supplementary materials. All materials are available through requests to the corresponding author.

## Acknowledgments

We thank Meirong Liang and IRCM animal facility staff for technical support; Drs. Jeffrey Mogil and Samantha Butler for comments on the manuscript; Dr. Farin Bourojeni and other members of the Kania laboratory for advice, feedback, and discussions. This work was supported by project grants from the Canadian Institutes of Health Research (CIHR; PJT-162225, MOP-77556, PJT-153053, PJT-159839, PJT-183824, and PJT-197987) to AK and National Institutes of Health (NIH; R01-NS050580) to LL. AK holds the Doggone Foundation Chair of Excellence in Pain and receives support from the Fondation IRCM. KTS is a recipient of a CIHR doctoral scholarship (FRN:494078). XZ is a recipient of a Fonds de recherche du Québec -Santé doctoral scholarship.

## Author contributions

K.T.S. Performed all experiments except for the in-vitro and behavioural experiments, analysed the data, and performed the computational modelling. The in-vitro experiments were performed and analysed by D.d.T. with reagents generated by E.S. Behavioural experiments using Ten3^KO^ mice were performed by A.M.M. and D.T.P. and quantified by S.M. Behavioural experiments using Ten3^ΔDH^ mice were performed by X.Z. and quantified by C.S. R.B.R contributed to investigation and conceptualization. L.L., S.P., T.B. and C.F.B. provided reagents. A.K. supervised the study and acquired funding. A.K. and K.T.S. wrote the paper.

---

**Figure S1.**
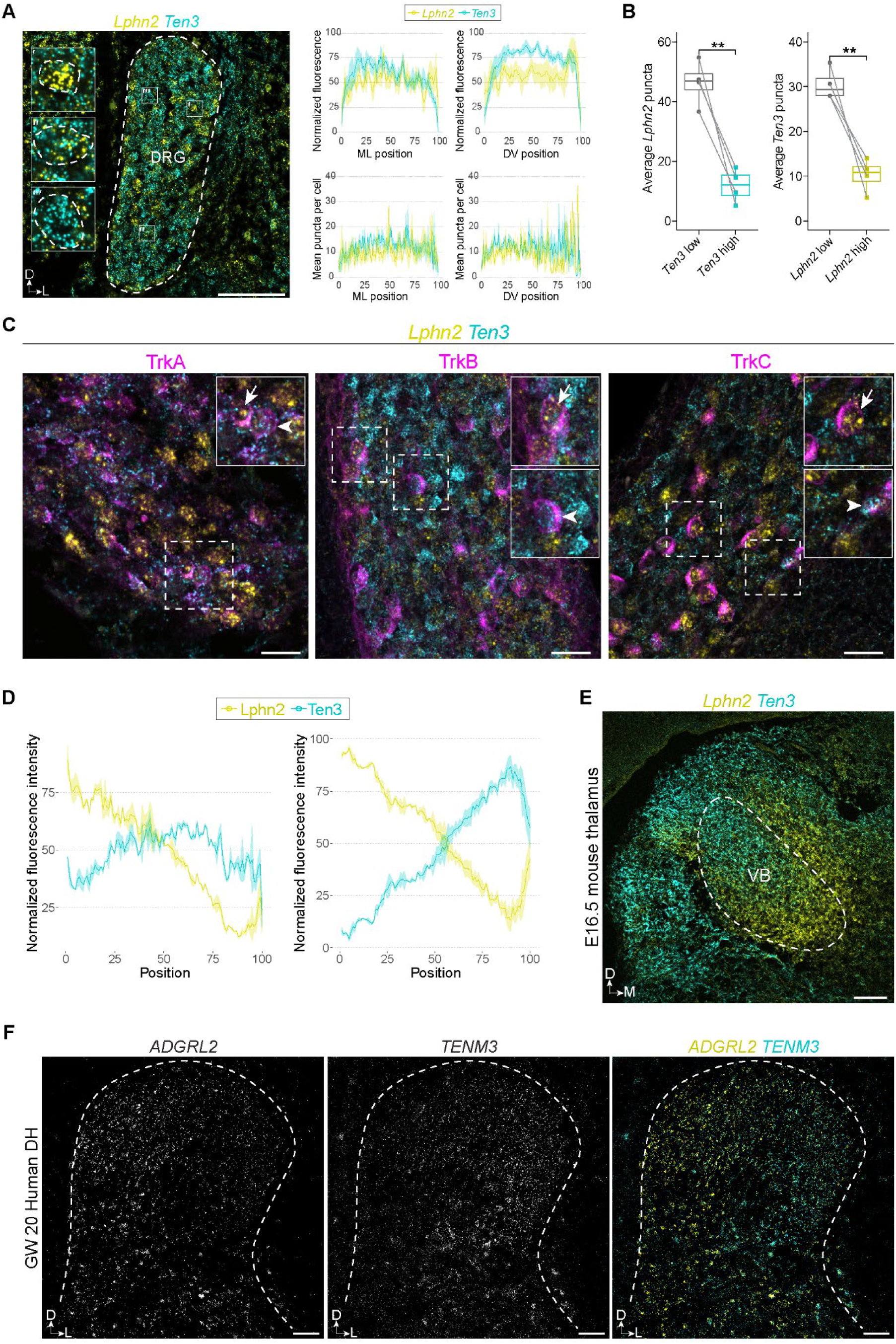
Further characterization of Ten3 and Lphn2 expression. **A**, *Ten3* and *Lphn2* mRNA localisation in e14.5 DRG cross sections (left) and quantification of puncta counts and fluorescence intensity (right, n = 4 embryos) showing no spatial bias for either across the DV or ML axes. Insets show DRG neurons with high *Lphn2* (′), a mix of *Lphn2* and *Ten3* (″), and high *Ten3* mRNA levels (′″). **B**, Quantification of mRNA puncta in (A) for *Lphn2* in *Ten3*-low compared to *Ten3*-high cells and or *Ten3* in *Lphn2*-low compared to *Lphn2*-high cells. High mRNA level cells were defined as the top 90% and low mRNA cells were defined as the bottom 10% of cells expressing each respective mRNA after filtering out cells with low/no expression of both. Counts per cell were averaged such that each point corresponds to one embryo (n = 4 embryos). **C**, Dual *Ten3* and *Lphn2* mRNA and TrkA, TrkB, and TrkC DRG neuron subtype protein marker detection in e14.5 DRGs. **D**, Related to Figure 1B, quantification of fluorescence intensity of immunostaining for Ten3 and Lphn2-mVenus fusion protein (detected with an anti-GFP antibody) in the white matter adjacent to the DH of *Lphn2-mVenus* knock-in mice at e14.5 (n = 3 embryos). Shown as fluorescence intensity normalized per image (left) or as a proportion of total fluorescence intensity at each ML position (right). **E**, *Ten3* and *Lphn2* mRNA detection in the e16.5 ventrobasal (VB) thalamus aligning with the proximal to distal somatotopic organization (dorsolateral to ventromedial VB). **F**, *TENM3* and *ADGRL2* (human orthologues of *Ten3* and *Lphn2*) mRNA detection in the human gestational week (GW) 20 DH. All data shown as mean ± SEM. A paired t-test was performed in **c**. ***P ≤ 0.001; **P ≤ 0.01; *P ≤ 0.05. Scale bars in **A** = 25 μm, in **B**, **E**, and **F** = 100 μm.

**Figure S2.**
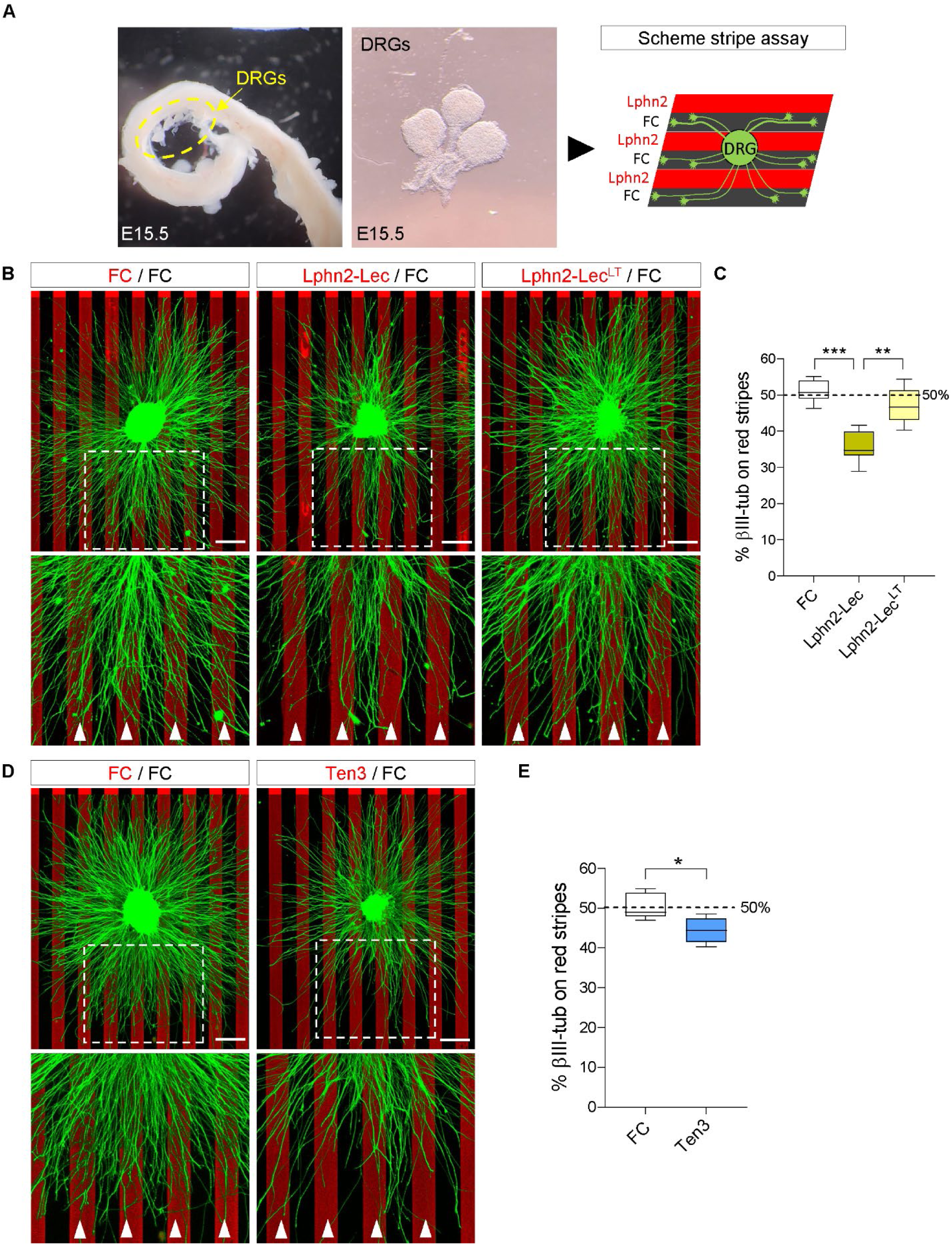
Ten3 and Lphn2 guide DRG axons *in vitro*. **A**, Schematic of DRG explant experiments. **B**, Explants of e15.5 lumbar DRGs labelled with anti-βIII-tubulin antibody, cultured on stripes of: FC (black), FC (red, left), Lphn2-lectin domain (Lphn2-Lec, red, middle), or Lphn2 non-Teneurin-binding mutants (Lphn2-Lec^LT^, red, right) proteins. Red stripes are also marked by white arrowheads. **C**, Quantification of βIII-tubulin protein on red stripes as a percentage of the total βIII-tubulin signal. While there is no preference for red or black stripes in either FC/FC or FC/ Lphn2-Lec^LT^ controls, there are significantly fewer βIII-tubulin labelled axons on the red Lphn2-Lec stripes. **D**, Explants of e15.5 lumbar DRGs labelled with anti-βIII-tubulin antibody grown on stripes of: FC (black), FC (red, left), or Ten3 (red, right) proteins. Red stripes are also marked by white arrowheads. **E**, Quantification of βIII-tubulin on red stripes as a percentage of the total βIII-tubulin signal. While there is no preference for red or black stripes in FC/FC controls, there are significantly fewer βIII-tubulin labelled axons on the red Ten3 stripes, indicating that despite the presumed presence of both Ten3**→**Lphn2 repulsion and Ten3**→**Ten3 attraction, the net effect is repulsion. ***P ≤ 0.001; **P ≤ 0.01; *P ≤ 0.05. Scale bars = 200 μm.

**Figure S3.**
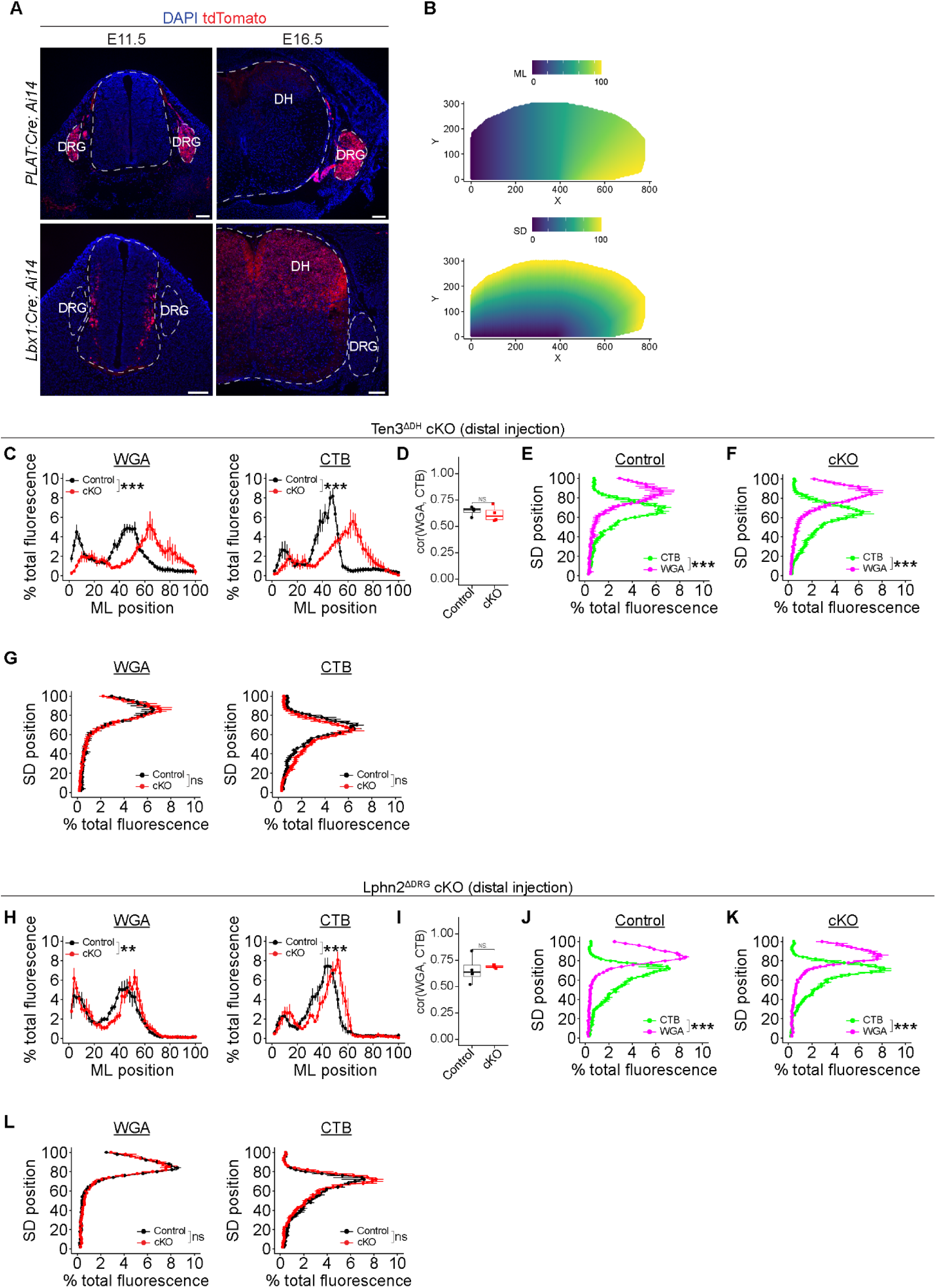
Additional characterization of distal sensory afferent termini in the DH of Ten3 and Lphn2 cKOs. (**A**) Validation of cre driver lines used in this study showing cre-driven tdTomato expression in the DH and DRG at E11.5 and E16.5 (**B**) A representative example showing mapping of x and y axes to anatomically-relevant axes used in quantifications in this study to account for DH curvature. (**C** and **H**) ML distribution of WGA (left) and CTB (right) in (C) Ten3^ΔDH^ cKOs and (H) Lphn2^ΔDRG^ cKOs compared to littermate controls. (**D** and **I**) Average Pearson correlation coefficient per animal between WGA and CTB signal across the ML axis of the DH in (D) Ten3^ΔDH^ cKOs and (I) Lphn2^ΔDRG^ cKOs compared to littermate controls. (**E**, **F**, **J** and **K**) SD distribution of CTB and WGA signal in controls (E and J) and cKOs (F and K) showing separation along the SD axis of afferents labeled by the two tracers. (**G** and **L**) SD axis distribution of WGA (left) and CTB (right) between cKOs and control littermates showing no shift along this axis. All data shown as mean ± SEM. Ten3^ΔDH^, n = 4 controls and 4 cKOs; Lphn2^ΔDRG^, n = 4 controls and 3 cKOs. Welch’s t-test was performed in (D) and (I), in all others two-way mixed ANOVA was performed and genotype x bin interaction term is reported. ***P ≤ 0.001; **P ≤ 0.01; *P ≤ 0.05. Scale bars = 100 μm.

**Figure S4.**
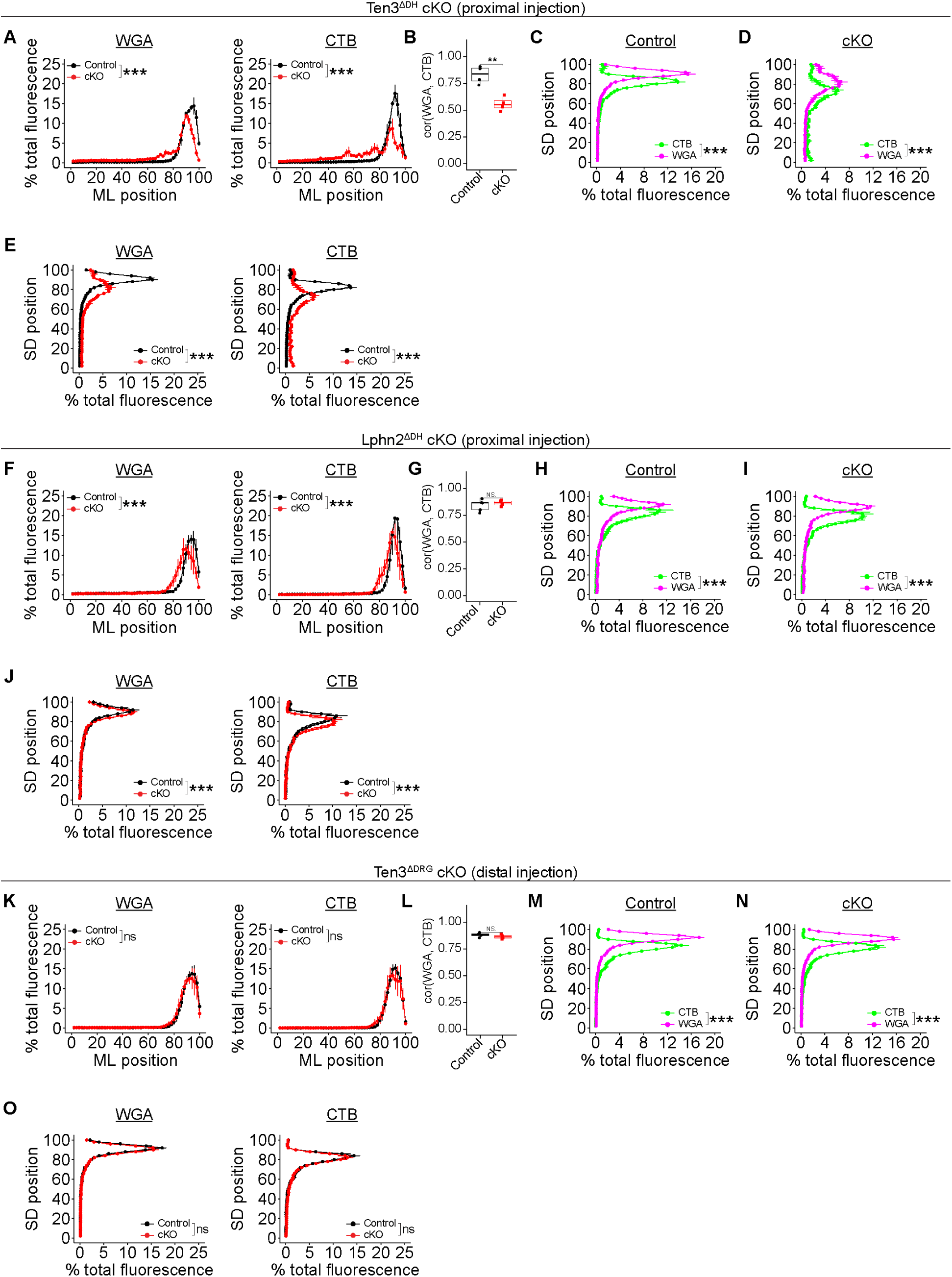
Additional characterization of proximal sensory afferents in the DH of Ten3 and Lphn2 cKOs. (**A**, **F** and **K**) ML distribution of WGA (left) and CTB (right) in (A) Ten3^ΔDH^ cKOs, (F) Lphn2^ΔDH^ cKOs, and (K) Ten3^ΔDRG^ cKOs compared to littermate controls. (**B**, **G** and **L**) Average Pearson correlation coefficient per animal between WGA and CTB signal across the ML axis of the DH in (B) Ten3^ΔDH^ cKOs, (G) Lphn2^ΔDH^ cKOs, and (L) Ten3^ΔDRG^ cKOs compared to littermate controls. (**C**, **D**, **H**, **I**, **M** and **N**) SD distribution of CTB and WGA signal in controls (C, H, and M) and cKOs (D, I, and N) showing separation along the SD axis of afferent termini labeled by the two tracers. (**E**, **J** and **O**) Distribution of WGA (left) and CTB (right) long the SD axis of cKOs and control littermates. All data shown as mean ± SEM. n = 4 controls and 4 cKOs for all analyses. Welch’s t-test was performed in (B), (G) and (L), in all others two-way mixed ANOVA was performed and genotype x bin interaction term is reported. ***P ≤ 0.001; **P ≤ 0.01; *P ≤ 0.05.

**Figure S5.**
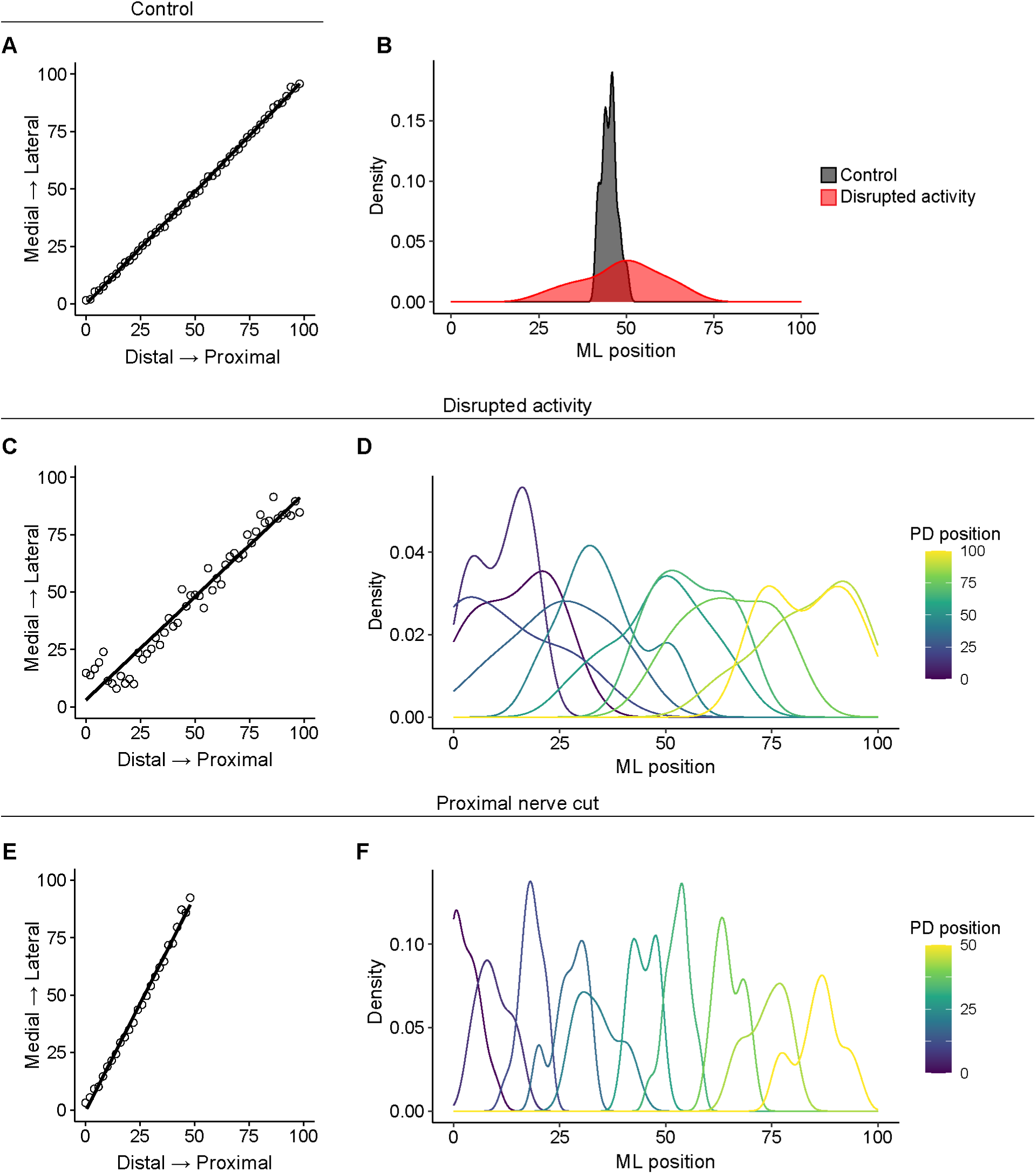
Further evaluation of computational model performance. (**A**) PD origin of DRG axons compared to the position of their ML terminal zones showing correct distal → medial and proximal → lateral mapping from one representative simulation under wild-type conditions. (**B**, **C**, and **D**) Simulated disruption of activity-dependent refinement from one representative simulation under wild-type conditions. (B) PD origin of DRG axons compared to the position of their ML terminal zones. (C) Simulated ML density of terminal zones in the DH from 10 DRG axons selected across the PD axis. (D) Comparison of one terminal zone from the same axon with/without disrupted activity-dependent refinement. (**E** and **F**) Simulated nerve cut experiment from one representative simulation under wild-type conditions showing expansion of remaining axons along the entire ML extent of the DH. (E) PD origin of DRG axons compared to the position of their ML terminal zones. (F) Simulated ML density of terminal zones in the DH from 10 DRG axons selected across the PD axis.

**Figure S6.**
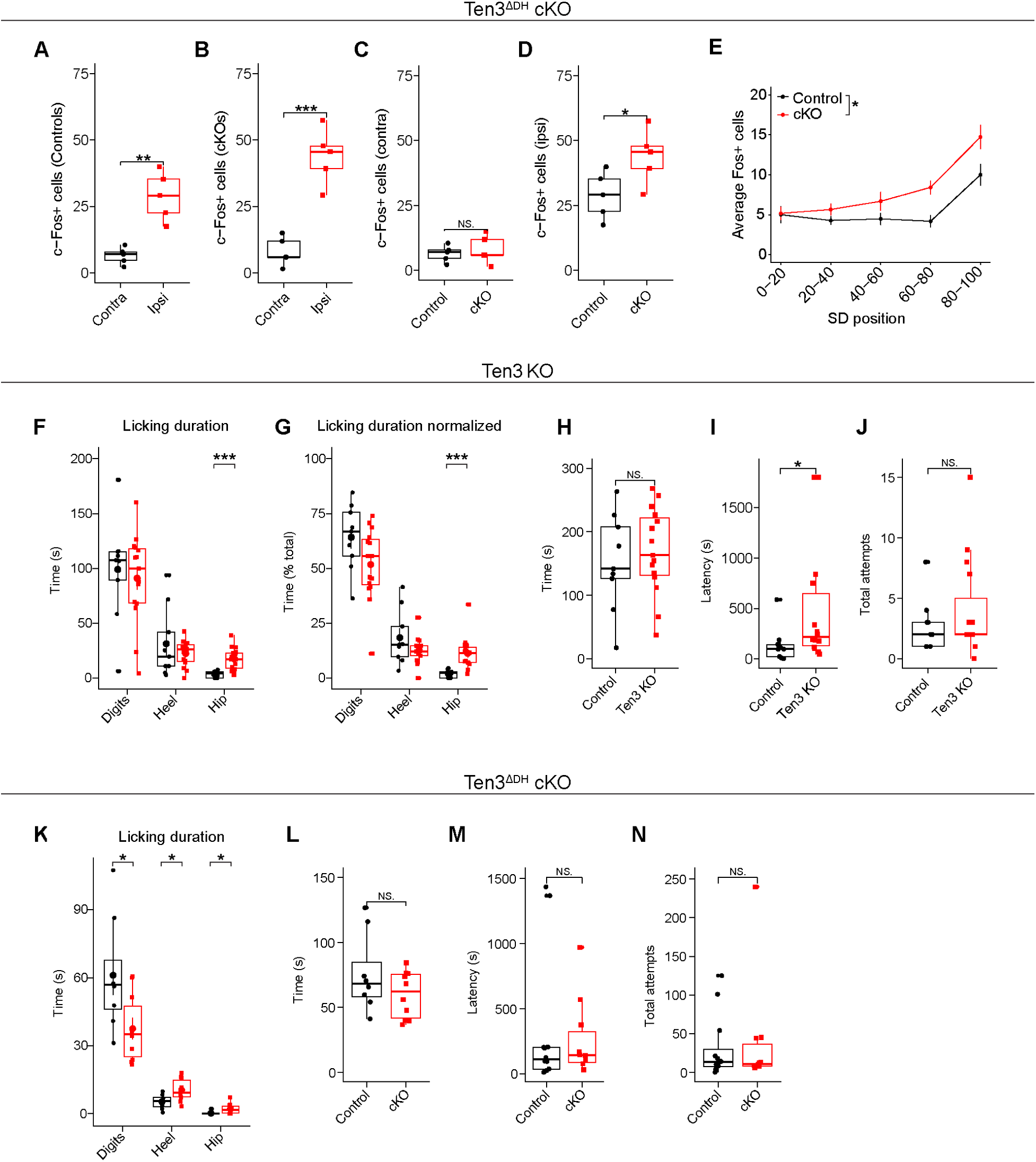
Additional characterization of c-Fos expression and somatosensory behavioural responses in Ten3 mutants. (**A** and **B**) Related to Figure 5, quantification of c-Fos-expressing cells in the contralateral and ipsilateral DH relative to the side of the formalin injection in (A) controls and (B) cKOs. (**C** and **D**) Quantification of c-Fos-expressing cells in cKOs compared to littermate controls in the (C) contralateral and (D) ipsilateral DH relative to the site of formalin injection. (**E**) SD distribution of c-Fos-expressing cells in cKOs compared to littermate controls in the ipsilateral DH. (**F-H**) Quantification of nocifensive licking behavior in Ten3 global KOs compared to control littermates in response to formalin injection to the hindpaw near the digits. (F) The percent of total time spent licking per limb region. (G) Licking time in seconds per limb region. (H) Total licking time. (**I**) Latency to remove adhesive tape from the hindpaw in KOs and control littermates. (**J**) Number of attempts to remove adhesive tape from the hindpaw in KOs and littermate controls. (**L-N**) Same as in panels (F), and (H-J) respectively, except performed in Ten3^ΔDH^ cKOs and control littermates. All data shown as mean ± SEM. (A-E) n = 5 controls and 5 cKOs; (F-H) n = 9 controls and 15 cKOs, (L-N) n = 8 controls and 8 cKOs. Welch’s t-test was performed in (A-D) and Mann-Whitney tests were performed in (H-J) and (K-N). In (F), (G) and (L) a two-way mixed ANOVA was performed, genotype x bin interaction term is reported, and post-hoc Mann-Whitney tests were performed for each limb location. ***P ≤ 0.001; **P ≤ 0.01; *P ≤ 0.05. Scale bars = 100 μm.

**Table S1.** List of computational model parameters. List of parameter values derived from PyABC along with their description and corresponding value in a retinocollicular model.

| Parameter category | Parameter | Value | Value in Triplet et al. 2011<sup>8</sup> | Description |
| --- | --- | --- | --- | --- |
| Activity-dependent refinement | $\gamma$ | 1.533263 | 0.05 | Overall strength of activity-dependent refinement |
| | $a_{act}$ | 3<br>(matched to Triplett et al.) | 3 | How the effect of a correlated activity scales with distance (in the target) |
| | $b_{act}$ | 11<br>(matched to Triplett et al.) | 11 | How the effect of a correlated activity scales with distance (in the periphery) |
| Competition | $A_{comp}$ | 683.569727 | 500 | Competition term (promotes more synapses) |
| | $B_{comp}$ | 0.410343 | 1 | Competition term (limits #synapses per axon) |
| | $D_{comp}$ | 3.152244 | 1 | Competition term (limits #synapses per dendrite) |
| Chemoaffinity | $\alpha_{chem}$ | 81.404452 | 20 | Overall strength of “chemoaffinity” (guidance cues) |
| | $k_{tl}$ | 0.522447 | NA | Strength of teneurin → latrophilin repulsion |
| | $k_{lt}$ | 0.789563 | NA | Strength of latrophilin → teneurin repulsion |
| | $k_{tt}$ | 0.226765 | NA | Strength of teneurin → teneurin attraction |
| Genotype/ activity dependent refinement interaction terms | $a_{t\_DH}$ | 2.991975 | NA | Slope of curve describing how loss of Ten3 in the DH should affect activity dependent refinement in each cell |
| | $b_{t\_DH}$ | 1.207797 | NA | Shape of curve describing how loss of Ten3 in the DH should affect activity dependent refinement in each cell |
| | $a_{t\_DRG}$ | 0.627226 | NA | Slope of curve describing how loss of Ten3 in the DRG should affect activity dependent refinement in each cell |
| | $b_{t\_DRG}$ | 1.532894 | NA | Shape of curve describing how loss of Ten3 in the DRG should affect activity dependent refinement in each cell |
| | $a_{l\_DH}$ | 1.505978 | NA | Slope of curve describing how loss of Lphn2 in the DH should affect activity dependent refinement in each cell |
| | $b_{l\_DH}$ | 1.250706 | NA | Shape of curve describing how loss of Lphn2 in the DH should affect activity dependent refinement in each cell |
| | $a_{l\_DRG}$ | 0.632682 | NA | Slope of curve describing how loss of Lphn2 in the DRG should affect activity dependent refinement in each cell |
| | $b_{l\_DRG}$ | 1.647495 | NA | Shape of curve describing how loss of Lphn2 in the DRG should affect activity dependent refinement in each cell |

